# Systematic characterization of a non-transgenic A*β*_1–42_ amyloidosis model: synaptic plasticity and memory deficits in female and male mice

**DOI:** 10.1101/2023.05.09.539973

**Authors:** Raquel Jiménez-Herrera, Ana Contreras, Guillermo Iborra-Lázaro, Danko Jeremic, Souhail Djebari, Juan Navarro-López, Lydia Jiménez-Díaz

**Affiliations:** Neurophysiology & Behavior Lab, Centro Regional de Investigaciones Biomédicas (CRIB), School of Medicine of Ciudad Real, University of Castilla-La Mancha, Ciudad Real, Spain

**Keywords:** Alzheimer’s disease, amyloid-*β*, spatial memory, LTP, hippocampus, sex differences.

## Abstract

**Background:** One of the neuropathological hallmarks of Alzheimer’s disease (AD) is amyloid-*β* (A*β*) accumulation in the hippocampus that causes its dysfunction. This disruption includes excitatory/inhibitory imbalance, synaptic plasticity and oscillatory activity impairments, and memory deficits. Although AD prevalence is higher in women than men, the possible sex difference is scarcely explored and information from amyloidosis transgenic mice models is contradictory. Thus, given the lack of data of the early amyloidosis stages in females, the aim of this study was to systematically characterize the effect of an intracerebroventricular (*icv.*) injection of A*β*_1–42_ on hippocampal-dependent memory, and on associated activity-dependent synaptic plasticity in the hippocampal CA1–CA3 synapse, in both male and female mice.

**Methods:** To do so, we evaluated long term potentiation (LTP) with *ex vivo* electrophysiological recordings and spatial (working, short- and long-term) and exploratory habituation memory using Barnes maze or open field habituation tasks respectively.

**Results:** We found that A*β*_1–42_ administration impairs all forms of memory evaluated, regardless the sex, in a long-lasting manner (up to 17 days post-injection). Furthermore, LTP was inhibited at a postsynaptic level, both in males and females, and a long-term depression (LTD) was induced for the same prolonged period, which could underly memory deficits.

**Conclusions:** In conclusion, our results provide further evidence of the shifting of LTP/LTD threshold due to a single *icv*. A*β*_1-42_ injection, which underly cognitive deficits in early stages of AD. These long-lasting cognitive and functional alterations in males and females validate this model for the study of early amyloidosis in both sexes, thus offering a solid alternative to the inconsistence of amyloidosis transgenic mice models.

## Background

The role of the hippocampus in learning and memory processes is well stablished (1, 2). Several types of memory, such as episodic memory, spatial memory, or contextual fear, are dependent on the hippocampus (3–5). Spatial memory is essential for the survival of all kind of animals, since it allows the retrieval of the location of an object, and to place an experience in a specific context in the environment (6). The encoding of spatial memory relies on the catching of behaviorally relevant spatial cues on a timescale of seconds, known as spatial working memory (7). Moreover, the hippocampus is one of the most vulnerable brain areas, prone to succumb to oxidative stress derived from different diseases (8). This can lead to the characteristic memory and learning impairments of pathologies like Alzheimer’s disease (AD), addiction or major depression, among others (9, 10).

AD is the main cause of dementia, affecting over 55 million people worldwide (11). One of the neuropathological hallmarks of AD is amyloid-*β* (A*β*) plaques accumulation in the hippocampus, known as senile plaques, seen preferentially in the late stages of the disease (12). A*β* causes a hippocampal dysfunction, which include excitatory/inhibitory (E/I) imbalance, impairments of hippocampal oscillatory activity, synaptic plasticity disruption and memory deficits (13–15). In the early stages of AD, electrophysiological data have proven that soluble A*β* oligomers can block hippocampal long-term potentiation (LTP) while enhancing long-term depression (LTD) (16, 17). These two mechanisms of synaptic strengthening and weakening, respectively, underlie synaptic plasticity, and the balance between both is highly important to maintain proper hippocampal functionality, such as learning and memory processes (18). Indeed, it has been previously demonstrated that LTP inhibition caused by A*β* correlates with the memory impairments reported in AD (15, 19–23).

As mentioned above, the accumulation of A*β* in the hippocampus is an important factor for AD development (16) found in both sporadic AD - the most common form of the disease- and familial inherited AD. Many studies use transgenic mice which express and accumulate the human A*β* peptide (Amyloid Precursor Protein-APP- models based on human mutations that represent less than 5% of AD cases (inherited AD)), *tau* hyperphosphorylation or both (24). However, the physiopathology of AD starts even decades before the senile plaques are present, when A*β* is soluble rather than accumulated (25). The A*β*_1–42_ fragment has higher propensity to form amyloid fibrils than other fragments and is found to be the dominant A*β* specie in the amyloid plaques of AD patients (24, 26). Moreover, it has been proven that intracerebroventricular (*icv.*) administration of A*β* diffuses mainly to the dorsal hippocampal formation (22), which is related specifically to spatial learning and memory (27, 28). Accordingly, this model seems a better option for investigating early stages of sporadic AD (29). In this line, previous work from our group have shown an E/I imbalance in male mice after an *icv.* injection of A*β*_1–42_ both *in vivo* and *in vitro*. This translated into a deficit in LTP induction, a disruption of the correct neural oscillatory synchronization in the hippocampus and, ultimately, an impairment in learning and memory processes (22, 23, 30).

According to the Alzheimer’s Association, nearly two-thirds of AD patients in the US are women (31). Even though AD prevalence is higher in women than men, almost all the available data comes from studies with male mice (32). Therefore, it is crucial to include both, male and female animals in this type of studies. A few recent works have done so, although the results are inconsistent, showing sex-dependent differences (20, 33–36) or not (37, 38) depending on the animal model used, the specific parameters measured and/or the experiments carried out.

Thus, given the lack of data regarding AD models in females, the aim of the present study was to characterize the effect of *icv.* A*β*_1–42_ administration on working, short-term and long-term memory, as well as on synaptic plasticity processes, to provide a solid model to study early stages of amyloidosis considering sex differences.

## Materials and methods

### Animals

Female and male C57BL/6 adult mice (12-24 weeks old; 20-30 g) were used (RRID:MGI:5,656,552; Charles River, US). Animals were kept on 12 h light/dark cycles with access to food and water *ad libitum*. Temperature (21 ± 1 °C) and humidity (50 ± 7%) were controlled. Mice were housed in same-sex groups of 5 per cage before surgery, and individually afterwards. All experimental procedures were carried out at the same time interval in both female and male mice to minimize circadian rhythm interferences.

All experimental procedures were reviewed and approved by the Ethical Committee for Use of Laboratory Animals of the University of Castilla-La Mancha (PR-2022-11-04 and PR-2018-05-11) and conducted according to the European Union guidelines (2010/63/EU) and the Spanish regulations for the use of laboratory animals in chronic experiments (RD 53/2013 on the care of experimental animals: BOE 08/02/2013).

### Surgery for drug injection

Mice were anesthetized with 4% isoflurane (#13400264, ISOFLO, Proyma S.L., Spain) using a calibrated R580S vaporizer (RWD Life Science; flow rate: 0.5 L/min O_2_). After induction, 1.5% isoflurane was delivered constantly for the maintenance of anesthesia. Intramuscular buprenorphine (0.01mg/kg; #062009, BUPRENODALE, Albet, Spain) and a healing cream (Blastoestimulina®; Almirall, Spain) were administered as analgesic after surgery, to accelerate recovery and decrease animal suffering.

As described elsewhere (23), for *icv*. administration of A*β*, animals were implanted with a blunted, stainless steel, 26-G guide cannula (Plastics One, US) in the left ventricle (1 mm lateral and 0.5 mm posterior to bregma; depth from brain surface, 1.8 mm) (39). The final position of the cannula was determined by Nissl staining (Figure 1A).

**Figure 1.**
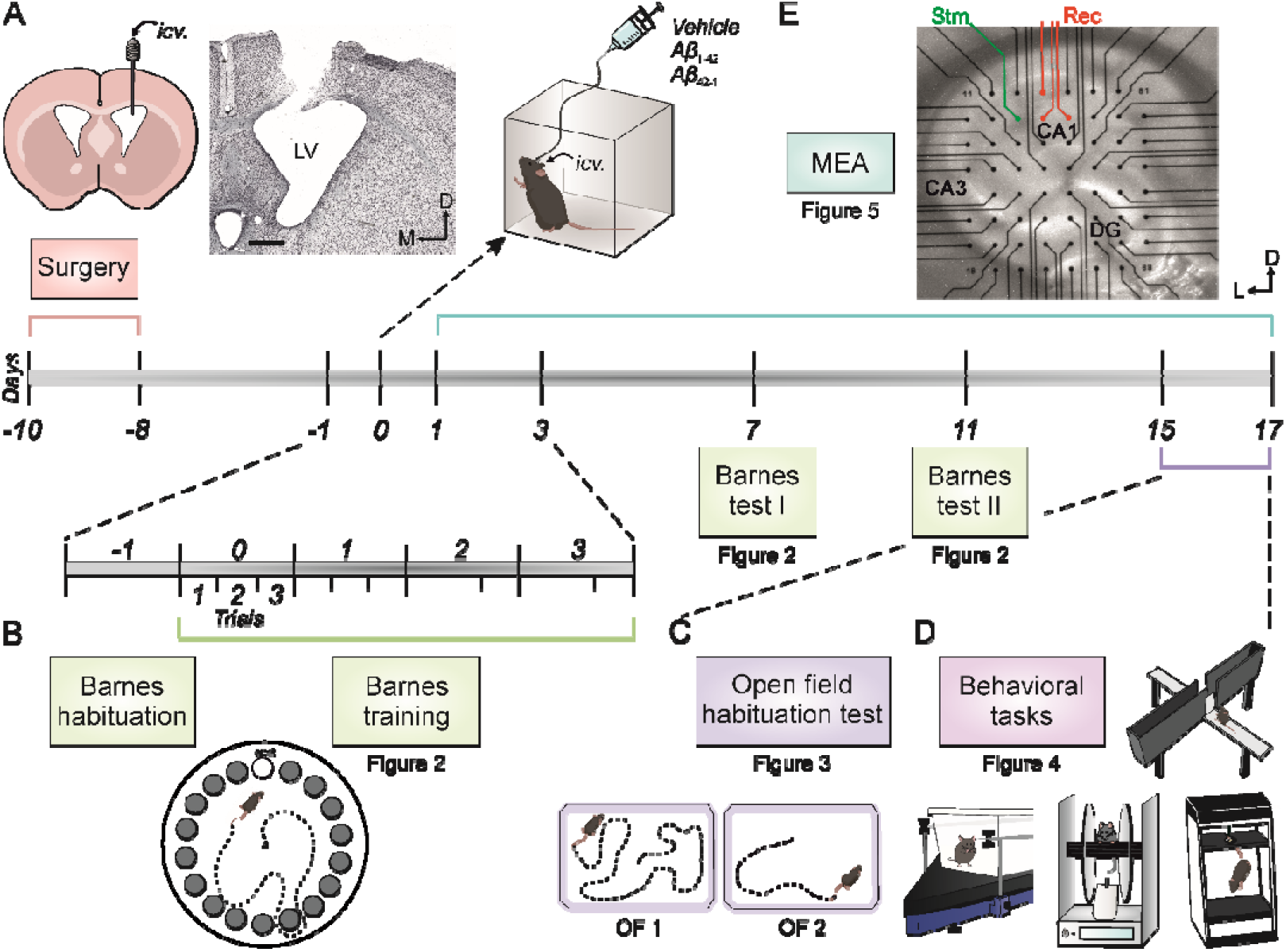
Experimental design showing the timeline and corresponding figure numbers. **(A)** A stainless-steel guide cannula was implanted for *icv*. drug administration on the left ventricle, as shown in the histologic verification image on the right. A minimum of 8 days after surgery (day 0), either A*β*_1-42_ or controls: reverse A*β*_42-1_ peptide or vehicle, were administered *icv.* Scale bars: 500 mm. **(B)** The Barnes maze task habituation phase was carried out the day before *icv.* administration (day -1). One hour after *icv*. injection of the corresponding treatment (day 0), training in the Barnes maze started. The training phase consisted of 3 trails per day, during 4 consecutive days (days 0 to 3). Two memory tests were applied, on days 7 and 11 post-injection. **(C)** The Open field habituation test was carried out on days 15 and 16 post injection. **(D)** Finally, mice went through a test battery to assess their overall spontaneous behavior and health, that consisted of testing of stereotyped and locomotor behaviors by an automated LABORAS^®^ System, as well as the rotarod test, the elevated plus maze and the tail suspension test (days 15 to 17). **(E)** Another cohort of animals was used to evaluate the effect of A*β*_1-42_ on *ex vivo* hippocampal LTP by multi-electrode arrays (MEAs) electrophysiology (days 1 to 17 post-injection). Representative location of stimulation (Stm, green) and recording (Rec, red) electrodes in a hippocampal coronal slice. D, dorsal; DG, dentate gyrus; *icv*., intracerebroventricular; L, lateral; LV, lateral ventricle; M, medial.

Mice were allowed at least a week for recovery before any experimental procedure. After recovery, freely moving animals received a 3 μg/µL *icv.* injection of either A*β*_1-42_, A*β*_42-1_ (as a reverse control) or vehicle (control, PBS) through an injection cannula at a rate of 0.5 μL/min. For this purpose, the injection cannula was inserted into the guide cannula protruding 0.5 mm into the ventricle and attached to a Hamilton syringe. All drugs were dissolved in PBS and purchased from Bachem (#4014447 and #4027991, respectively; Switzerland). Mice were routinely handled to minimize stress during experimental procedures.

### Barnes maze

To evaluate hippocampal-dependent working, short-term and long-term spatial memory, the Barnes maze (LE851BSW, Panlab, Spain) was used (40). It consists of a rotating platform disk (92 cm in diameter) with 20 scape holes (5 cm each) around its periphery, elevated 1 meter above the floor. Spatial cues (circles and squares of different colors) and mildly aversive white noise were used.

The protocol used was adapted from Suarez et al. (41). Briefly, each trial began with the mouse inside a starting cylinder (8 x 12.5 cm) positioned in the center of the maze. After 10 sec, the starting cylinder was removed, and the white noise was initiated. Between each trial, the maze was cleaned with 70% ethanol to dissipate odor cues. As shown in Figure 1B, the protocol consisted of a habituation trial the day before *icv.* injection (Day−1), four training days of three trials each, starting the day of treatment administration (Days 0 - 3), and two memory tests (Days 7 and 11). During the habituation day, mice were able to explore the maze for 90 sec or until they found the goal box (17.5 x 7.5 x 8 cm) attached to one of the holes. If time ran out, the animal was gently led to the goal box. In both cases, mice were kept in the box for 1 min before returning them to their cages. Afterwards, training was carried out on four consecutive days, with three trials per day and a 15 min interval between trials. During each trial, mice were allowed to explore the maze for 3 min or until they found the escape hole, which was changed daily. Latency to find the box, errors made, and distance traveled were measured. Finally, a single 90 sec trial was performed each memory testing day, without any scape hole available. Latency and distance to the target of the latest training day were measured. All sessions were recorded and analyzed with Barnes-Smart video tracking software (Panlab, Spain).

### Open field habituation task

On days 15 and 16 post-*icv.* injection (Figure 1C), to evaluate a non-associative hippocampal-dependent learning process, such as exploratory habituation to a novel environment (42), an open field (OF) habituation task was conducted as described elsewhere (43). Briefly, mice were exposed twice during consecutive days to an OF, to measure the change in exploration after re-exposure. In the training day (OF1), mice were placed in the middle of a square acrylic box (23.5 x 17.5 x 4 cm plexiglas base arena; 26.5 x 21 x 10 cm top) and allowed to freely explore it for 15 min. In the retention day (OF2), 24 h later, mice were re-exposed to the same environment. Exploratory behavior was recorded using a LABORAS^®^ apparatus (*Laboratory Animal Behavior Observation Registration and Analysis System*; Metris, Netherlands), which transforms mechanical vibrations generated by the movements of the animals into electrical signals through a sensing platform positioned under the cage.

### Spontaneous behaviors

Between days 15 and 17 post-injection, mice went through a battery of behavioral tests to assess their overall state and spontaneous behaviors (Figure 1D).

### Laboratory animal behavior observation registration and analysis system (LABORAS^®^) for stereotyped and locomotion behavioral testing

Mice were placed in a rectangular LABORAS^®^ cage for a single 15-min trial, as previously described (43). Grooming behavior was used as a measurement of stress-related behavior (44). Locomotion, climbing, and rearing behaviors were used as measurements of locomotor activity (45). All data were digitized and analyzed using the LABORAS^®^ software (Metris, Netherlands).

### Rotarod performance test

The rotarod apparatus (LE 8500, Panlab, Spain), consisting of a 30 mm diameter black striated rod positioned 20 cm above the floor, was used to measure coordination and motor function. Mice were trained to stay for 1 min on the rod at constant low-speed (6 rpm) rotation. The following day, 5 consecutive trials were carried out, in which the rod was set to accelerate from 4 to 40 rpm for 2 min, and mice’s time to fall off was recorded.

### Elevated plus maze

To assess anxiety-like behaviors (46), the elevated plus maze (LE 842, Panlab, Spain) was used. It consists of a cross-shaped methacrylate platform with two open arms (65 x 6 cm) without walls and two arms enclosed (65 x 6 cm) by 15-cm-high opaque walls, mounted 90° to one another with a central platform (6.3 x 6.3 cm) and raised 40 cm above the floor. Mice were placed into one of the open arms and were allowed to freely explore the maze during a single 5 min session. The number of entries into the open arms and the time spent there were calculated to measure anxiety-like behavior. Additionally, as a measurement of locomotor activity (47), number of entries into the closed arms and total entries (open + closed arms) were recorded.

### Tail suspension test

To assess depression-like behaviors, the tail suspension test was performed (48). The tail suspension apparatus (BIO-TST5, Bioseb, US) consists of three PVC chambers (50 x 15 x 30 cm each) in which animals were hung by the tail for 6 min approximately 10 cm away from the ground. The immobility time was registered by the strain sensors. All data were digitized and analyzed using the BIO-TST5 software (Bioseb, US).

### *Ex vivo* field EPSP (fEPSP) recordings

Coronal hippocampal slices were prepared as described previously (43). In brief, animals were deeply anesthetized with halothane (Fluothane, AstraZeneca, UK) and decapitated. The brain was immediately removed and rapidly immersed in oxygenated (95% O_2_ – 5% CO_2_) ice-cold “cutting” solution containing (in mmol/L; all from Sigma, US): 87 NaCl (#S9888), 10 glucose (#G8270), 75 sucrose (#84100), 1.25 NaH_2_PO_4_ (#S8282), 3 C_3_H_3_NaO_3_ (#P2256), 0.98 C_6_H_7_NaO_6_ (#11140), 25 NaHCO_3_ (#S6014), 2.5 KCl (#P3911), 0.37 CaCl_2_ (#499609), 3.28 MgCl_2_ (#208337). Brain was trimmed and mounted on the stage of a vibratome (7000smz-2; Campden Instruments, UK) in such a way so that the blade would cut through hemispheres at an angle of 20-30° of their horizontal planes. Coronal slices (300 µm thick) containing the dorsal hippocampus were then incubated for at least 1.5 h before recording, at room temperature (22°C) in oxygenated artificial cerebrospinal fluid (aCSF) containing (in mmol/L; all from Sigma, US): 125.99 NaCl (#S9888), 3 KCl (#P3911), 1.8 CaCl_2_ (#499609), 1.5 MgCl_2_ (#208337), 25.99 NaHCO_3_ (#S6014), 10 glucose (#G8270), and 1.2 NaH_2_PO_4_ (#S8282).

For electrophysiological recordings, two set-ups consisting of a multi-electrode array (MEA2100-Mini-System) pre-amplifier and a filter amplifier (gain 1100× or 550×) were run in parallel by a data acquisition card governed by MC_Experimenter V2.20.0 software. A single slice was transferred to each MEA recording chamber (MEA60; Multi Channel Systems, Reutlingen, Germany) continually perfused with aCSF (flow rate 2 mL/min) and kept at 32 °C. The MEA was positioned on the platform of an inverted MEA-VMTC-1 video microscope and it comprised 60 extracellular electrodes (inter-electrode distance: 200 μm). Each individual electrode from the array could be used either as a recording or as a stimulation electrode. A nylon mesh was positioned above the slice to obtain a satisfactory electrical contact between the surface of the slice and the electrode array. Stimulation was achieved with a stimulus generator unit integrated in the headstage (Multi Channel Systems, Germany) by applying biphasic current pulses to one electrode of the array (S1) located in the Schaffer Collateral pathway of the hippocampus. Field excitatory postsynaptic potentials (fEPSPs) could then be recorded in the *stratum radiatum* of the CA1 subfield by all the remaining electrodes of the array at the same time (Figure 1E). A second electrode of the array (S2) was used to stimulate an independent pathway without High Frequency Stimulation (HFS, stimulation protocol used to induce LTP; see details below), as control of synaptic transmission.

Following at least 20 min of equilibration period inside the MEA well, basal synaptic transmission was studied by input/output (I/O) curves, by applying two stimuli of increasing intensities (0.02 – 0.4 mA) at 40 ms interstimulus interval. After I/O, pulses were adjusted to ≈40% of the intensity necessary to evoke a maximum fEPSP response. To address a typical short-term plasticity phenomenon at a presynaptic level, the paired-pulse facilitation (PPF) protocol was used. Thus, pairs of stimuli were delivered at different interstimulus intervals (10, 20, 40, 100, 200, 500 ms). For LTP induction, a high-frequency stimulation (HFS) protocol was used, consisting of five 1-sec-long 100-Hz trains delivered at 30 sec intertrain interval. Baseline (BL) values of fEPSPs amplitude recorded at the CA3-CA1 synapse were collected for 15 min before LTP induction. After HFS, fEPSPs were recorded during 60 min to evaluate LTP induction. To pair it with the behavioral tasks, electrophysiology was carried out 1 to 17 days post-*icv.* injection of A*β*_1-42_ or controls: A*β*_42-1_ and vehicle.

Data was analyzed with the Multichannel Analyzer software (V 2.20.0). As synaptic responses were not contaminated by population spikes, the amplitude (i.e., the peak-to-peak value in mV during the rise-time period) of successively evoked fEPSPs was used. All values represented mean ± SEM with *n* indicating number of slices. Within each slice, data from 3 different recording electrodes was used.

### Statistical analysis

Data was represented as the mean ± SEM and analyzed by two-way ANOVA, followed by Tukey’s *post-hoc* analysis. When comparing only two groups, Student t test was used. Statistical significance was set at *p* < 0.05. All analyses were performed using SPSS software v.24 (RRID:SCR_002865; IBM, US) and GraphPad Prism software v.8.3.1 (RRID:SCR_002798; Dotmatics, US). Final figures were prepared using CorelDraw X8 Software (RRID:SCR_014235; Corel Corporation, Canada).

## Results

Firstly, we wondered whether *icv.* injection of A*β*_1-42_ would affect spatial and habituation learning and memory in a sex-specific manner. Thus, female and male mice carried out a protocol to measure spatial learning and memory using a Barnes maze, as well as an exploratory habituation protocol using an OF to assess this type of non-associative memory.

### A*β*_1-42_ impairs spatial learning and memory in both female and male mice

For the Barnes Maze task, animals performed 3 trails per day during 4 consecutive days (days 0-3), starting one hour after *icv.* injections (performed in day 0, Figure 1) of either A*β*_1-42_ or both controls: vehicle and reverse A*β*_42-1_ peptide. Differences in the latency to find the open hole between the first and last trial of each day (Figure 2A) is considered an assessment of working memory (41). No differences between males and females within each treatment group were found. Nevertheless, our data showed a significant treatment (Male: F_(2,33)_ = 7.755, *p* = 0.0017; Female: F_(2,35)_ = 8.719, *p* < 0.0008) and time (Male: F_(5.584,169.9)_ = 13.43, *p* < 0.001, Geisser-Greenhouse’s correction; Female: F_(5.706,185)_ = 8.838, *p* < 0.001, Geisser-Greenhouse’s correction) effect when comparing the latency to find the escape hole in the first and last trial from each training session, proving that both vehicle (male, n = 14; female, n = 15) and A*β*_42-1_ (male, n = 7; female, n = 6) groups had a normal spatial working memory while A*β*_1-42_ animals (male, n = 14; female, n = 16) showed a deterioration of this kind of memory, regardless of sex. Furthermore, the evolution of the different parameters measured along the four training days allowed us to evaluate short-term memory, since animals should remember how to perform the task and therefore latencies would decrease over time, even if the target hole was changed. Accordingly, our data showed a treatment effect for latency (Figure 2B; Male: F_(2,124)_ = 47.97, *p* < 0.001; Female: F_(2,128)_ = 23.12, *p* < 0.001), number of errors (Figure 2C; Male: F_(2,33)_ = 18.14, *p* < 0.001; Female: F_(2,127)_ = 12.09, *p* < 0.001) and distance traveled (Figure 2D; Male: F_(2,34)_ = 4.59, *p* = 0.017; Female: F_(2,35)_ = 11.30, *p* = 0.0002). *Post-hoc* analysis revealed that A*β*_1-42_ disrupted spatial learning in both female and male animals. Finally, to study long-term memory, two memory tests were conducted in days 7 and 11 post-injection, during which all holes were closed (Figure 2E) and latency to reach the target hole of the last training day (day 3), as well as distance traveled, were quantified. Our results showed a statistically significant difference in latency (Figure 2F; Male: F_(2,62)_ = 7.644, *p* = 0.0011; Female: F_(2,62)_ = 9.618, *p* < 0.001) without affecting distance traveled (Figure 2G). Once again, *post-hoc* analysis revealed that the differences were due to a worst performance of A*β*_1-42_-treated mice. Thus, these data suggest an impairment in hippocampal-dependent spatial working, short- and long-term memory and learning processes induced by A*β*_1-42_ in both sexes. In contrast, same concentrations of the control reverse peptide, A*β*_42-1_, did not differ from the control vehicle suggesting that the decline induced by A*β*_1-42_ was therefore specific.

**Figure 2.**
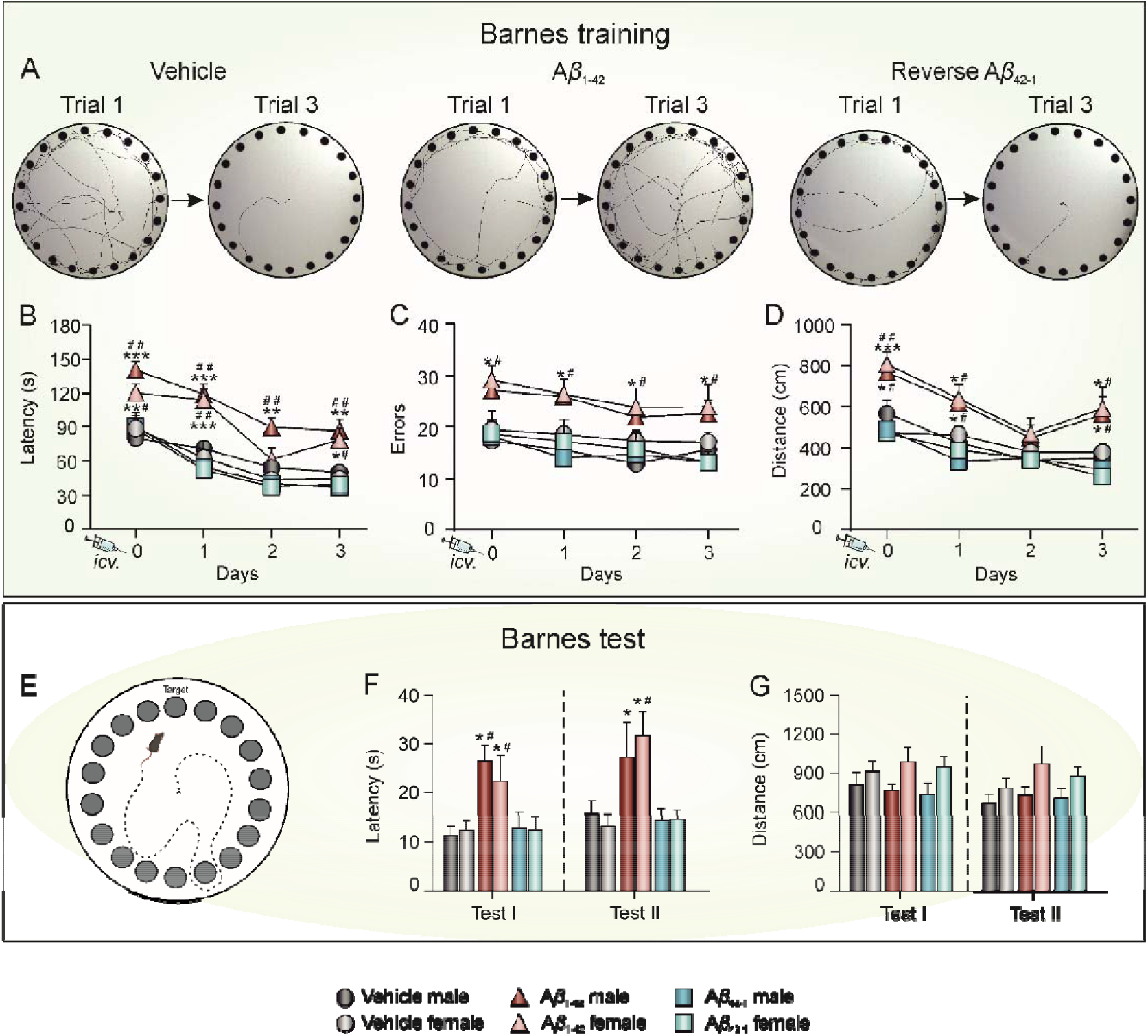
A*β*_1-42_ impairs spatial learning and memory in both female and male mice. **(A)** Representative traces of the path traveled during the first and last trial of the last training day (day 3) for A*β*_1-42_ and controls: vehicle and reverse A*β*_42-1_ peptide. **(B-D)** Escape latency (**B**; in s), number of errors (**C**) and distance traveled (**D**; in cm) during the four training days. **(E)** Overview image of the test phase on the Barnes maze, with all holes closed. **(F)** Latency (in s) to reach the target hole of the latest training day for the first time, during the two test sessions. **(G)** Distance traveled (in cm) during the two test sessions. Data is expressed as the mean ± SEM of the 3 trials per day. A*β*, Amyloid-*β*; cm, centimeters; s, seconds. * *p* < 0.05, ** *p* < 0.01, *** *p* < 0.001 *vs*. vehicle of the corresponding sex; # *p* < 0.05, ## *p* < 0.01 *vs*. A*β*_42-1_ of the corresponding sex.

### A*β*_1-42_ impairs exploratory habituation memory in both female and male mice

Afterwards, exploratory habituation memory was tested in days 15 and 16 post-*icv.* injection by an OF habituation task (Figure 3A). Data showed no significant differences in exploration between groups during the training session (OF1). However, when comparing OF1 and the retention session (OF2) (Figure 3B), a significant decrease in exploration was observed in both controls, vehicle (Male: *t*_12_ = 2.317, *p* = 0.019, n = 7; Female: *t*_18_ = 4.751, *p* < 0.001, n = 12) and A*β*_42-1_ (Male: *t*_12_ = 6.195, *p* < 0.001, n = 7; Female: *t*_10_ = 6.433, *p* < 0.001, n = 6) treated animals, regardless of sex, proving that they could remember the arena, while A*β*_1-42_ mice (male, n = 9; female, n = 10) performed slightly better without reaching significance, indicating some deterioration of memory in this group of animals. Moreover, during the retention session (OF2), a significant treatment effect was found in both female (Figure 3B; F_(2,22)_ = 4.450, *p* = 0.024) and male (Figure 3B; F_(2,20)_ = 4.952, *p* = 0.018) mice. *Post-hoc* analyses revealed that A*β*_1-42_ mice traveled a longer distance during this session than vehicle and A*β*_42-1_ control groups, as illustrated in Figure 3C. Hence, both results show that A*β*_1-42_ also impairs non-associative habituation memory even 2 weeks after treatment.

**Figure 3.**
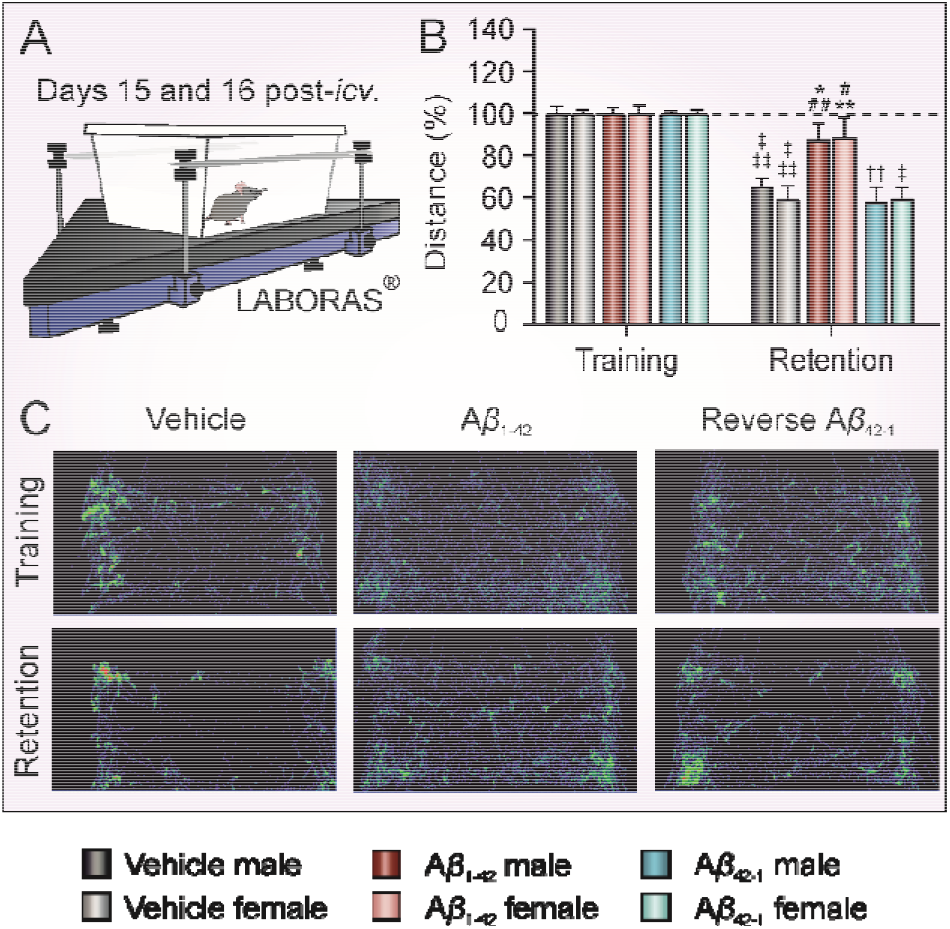
A*β*_1-42_ administration similarly alters non-associative exploratory habituation memory in both female and male mice. **(A)** An OF habituation test was carried out by submitting the animals to the same OF arena twice, on consecutive days 15 and 16 post-*icv.* injection. **(B)** Total distance traveled during the two OF sessions (training -OF1- and retention -OF2- sessions). Data is expressed as the percentage (%) of the distance traveled in the training session (OF1). **(C)** Examples of mice movement tracked during OF1 and OF2 for the different treatment groups. A*β*, Amyloid-*β*; *icv*., intracerebroventricular; OF, open field. * *p* < 0.05, ** *p* < 0.01 *vs*. vehicle of the corresponding sex; # *p* < 0.05, ## *p* < 0.01 *vs*. A*β*_42-1_ of the corresponding sex; ‡ *p* < 0.05, ‡‡ *p* < 0.01, ‡‡‡ *p* < 0.001 *vs*. OF1.

### Memory impairments were not due to health or locomotor disfunction

To verify that the deterioration observed after A*β*_1-42_ injection both in hippocampal-dependent spatial and habituation memory were due to specific hippocampal alterations, and rule out general health affections, mice carried out a battery of tests to evaluate different behavioral aspects. When assessing stereotyped behaviors using the LABORAS®, data showed that all groups (vehicle male, n = 11; vehicle female, n = 12; A*β*_1-42_ male, n = 11; A*β*_1-42_ female, n = 12; A*β*_42-1_ male, n = 7; A*β*_42-1_ female, n = 6) spent the same amount of time performing the different behaviors analyzed (Figure 4A): locomotion (Male: F_(2,28)_ = 0.208, *p* = 0.814; Female: F_(2,29)_ = 0.493, *p* = 0.616), climbing (Male: F_(2,28)_ = 0.596, *p* = 0.558; Female: F_(2,29)_ = 1.027, *p* = 0.372), rearing (Male: F_(2,28)_ = 0.962, *p* = 0.395; Female: F_(2,29)_ = 0.434, *p* = 0.653) and grooming (Male: F_(2,27)_ = 0.850, *p* = 0.439; Female: F_(2,29)_ = 1.120, *p* = 0.341). All groups equally improved their performance in the rotarod (Figure 4B; vehicle male, n = 8; vehicle female, n = 11; A*β*_1-42_ male, n = 8; A*β*_1-42_ female, n = 9; A*β*_42-1_ male, n = 7; A*β*_42-1_ female, n = 6), as no differences in the latency to fall off the rod were found between groups along trials (Male: F_(2,20)_ = 0.4095, *p* = 0.6694; Female: F_(2,23)_ = 0.09378, *p* = 0.9108) nor in the whole session (Male: F_(2,20)_ = 1.783, *p* = 0.1939; Female: F_(2,23)_ = 0.2739, *p* = 0.7629). Locomotion was also tested in all the groups (vehicle male, n = 8; vehicle female, n = 12; A*β*_1-42_ male, n = 9; A*β*_1-42_ female, n = 12; A*β*_42-1_ male, n = 7; A*β*_42-1_ female, n = 6) by the number of entries into closed (Male: F_(2,23)_ = 0.951, *p* = 0.403; Female: F_(2,29)_ = 0.116, *p* = 0.891) and total arms (Male: F_(2,23)_ = 0.255, *p* = 0.777; Female: F_(2,29)_ = 0.293, *p* = 0.748) in the elevated plus maze and, once again, no differences were found (Figure 4C). Regarding behaviors related to stress, all animals had the same number of entries (Figure 4C; Male: F_(2,23)_ = 0.663, *p* = 0.526; Female: F_(2,29)_ = 0.381, *p* = 0.687) and time spent in open arms (Male: F_(2,23)_ = 0.273, *p* = 0.763; Female: F_(2,29)_ = 0.391, *p* = 0.680) in the elevated plus maze, and they all had the same immobility time (Figure 4D; vehicle male, n = 8; vehicle female, n = 12; A*β*_1-42_ male, n = 9; A*β*_1-42_ female, n = 12; A*β*_42-1_ male, n = 7; A*β*_42-1_ female, n = 6; Male: F_(2,23)_ = 0.003, *p* = 0.997; Female: F_(2,29)_ = 0.077, *p* = 0.927) in the tail suspension test, suggesting that the treatment did not increase depression- nor anxiety-like behaviors.

**Figure 4.**
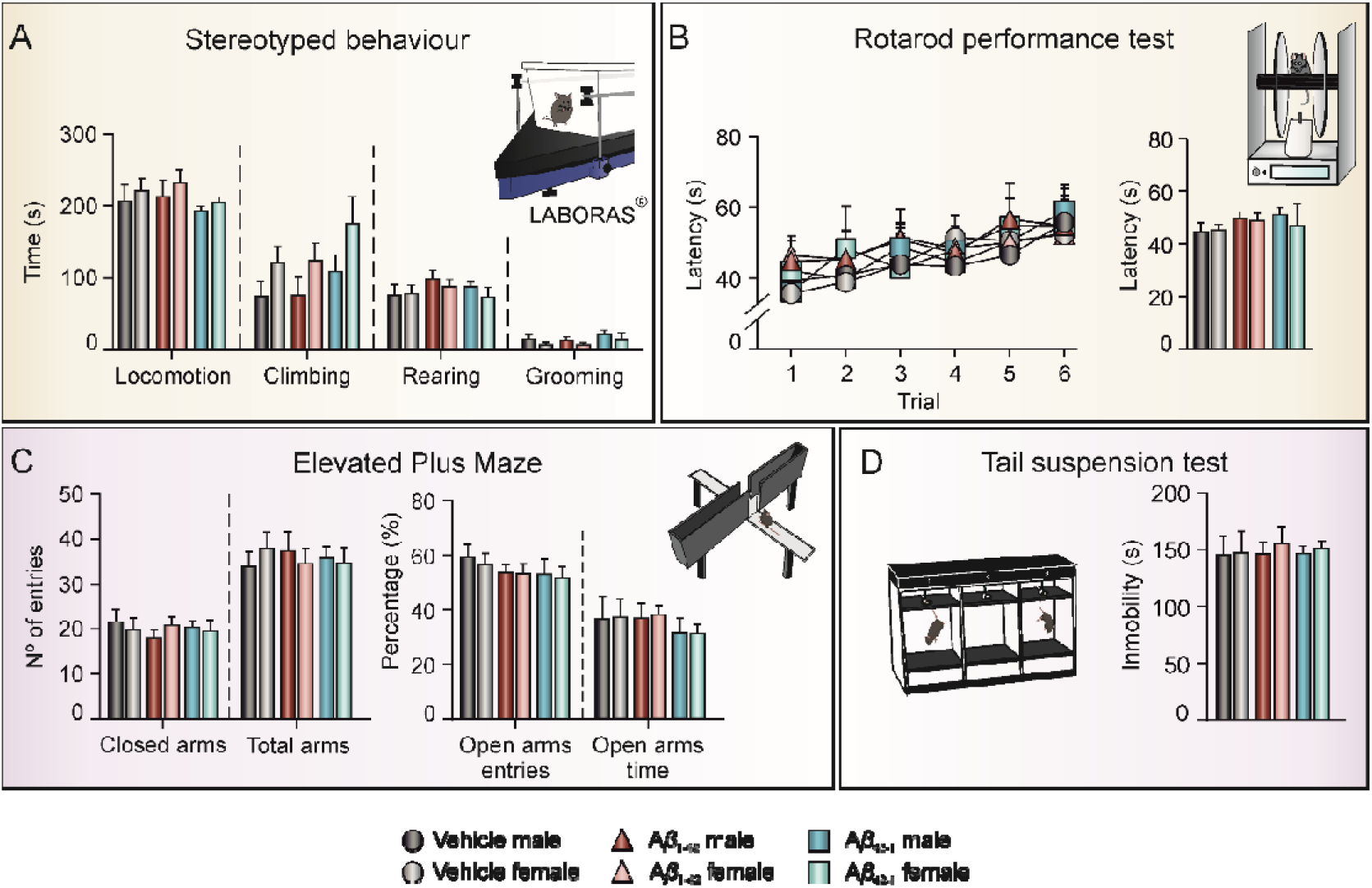
A*β*_1-42_ administration does not induce alterations in locomotor activity, anxiety, and depression-like behavior. Behavioral tasks to evaluate general health state were carried out on days 15 - 17 post-*icv.* injection. **(A)** Stereotyped behaviors were assessed using a LABORAS^®^ system, measuring the time (in s) spent performing each type of activity (locomotion, climbing, rearing, and grooming). **(B)** Latency (in s) to fall off the Rotarod during the six trials (left) and the whole session (right). **(C)** Number of entries in closed and total arms were used as measure of locomotor activity (left), while anxiety levels were assessed by the percentage (%) of entries and time spent on open arms in an elevated plus maze (right). **(D)** Depression-like behavior was assessed by measuring the immobility time (in s) during a single session in the tail suspension test. A*β*, Amyloid-*β*; s, seconds.

Thus, this data confirmed that the overall health state and locomotor function was uniform among the different treated groups and, therefore, all learning and memory impairments observed were due to a specific hippocampal disruption caused by A*β*_1-42_.

### A*β*_1-42_ inhibits *ex vivo* LTP in both female and male mice similarly

Given the deleterious effects of A*β*_1-42_ on hippocampal-dependent learning and memory processes in the present amyloidosis model in a similar way regardless the sex, we wondered whether excitability, presynaptic function, and short- and long-term plasticity were affected, since they are the underlying physiological mechanisms of those cognitive capabilities. To pair it with the behavioral tasks, electrophysiology was carried out 1 to 17 days post-*icv.* injection of A*β*_1-42_ or controls: A*β*_42-1_ and vehicle in a new cohort of mice.

Firstly, I/O curves in all groups (Figures 5A-C) showed a greater amplitude of both, the first (vehicle male, n = 7; vehicle female, n = 7; A*β*_1-42_ male, n = 8; A*β*_1-42_ female, n = 5; A*β*_42-1_ male, n = 5; A*β*_42-1_ female, n = 4; Male: F_(2.18,159.2)_ = 462.3, *p* < 0.001; Female: F_(2.28,157.3)_ = 470.1, *p* < 0.001, Geisser-Greenhouse’s correction) and the second fEPSP (Male: F_(2.221,126.6)_ = 242.2, *p* < 0.001; Female: F_(2.259,81.31)_ = 234.6, *p* < 0.001, Geisser-Greenhouse’s correction) with increasing intensities. No differences between groups were observed for either 1^st^ or 2^nd^ fEPSP in female slices, however in male slices a treatment effect was found in fEPSP2 (F_(2,57)_ = 4.297, *p* = 0.0183) since amplitudes evoked by the second pulse in A*β*_1-42_ injected animals were higher than those evoked in the other sex-matched groups.

**Figure 5.**
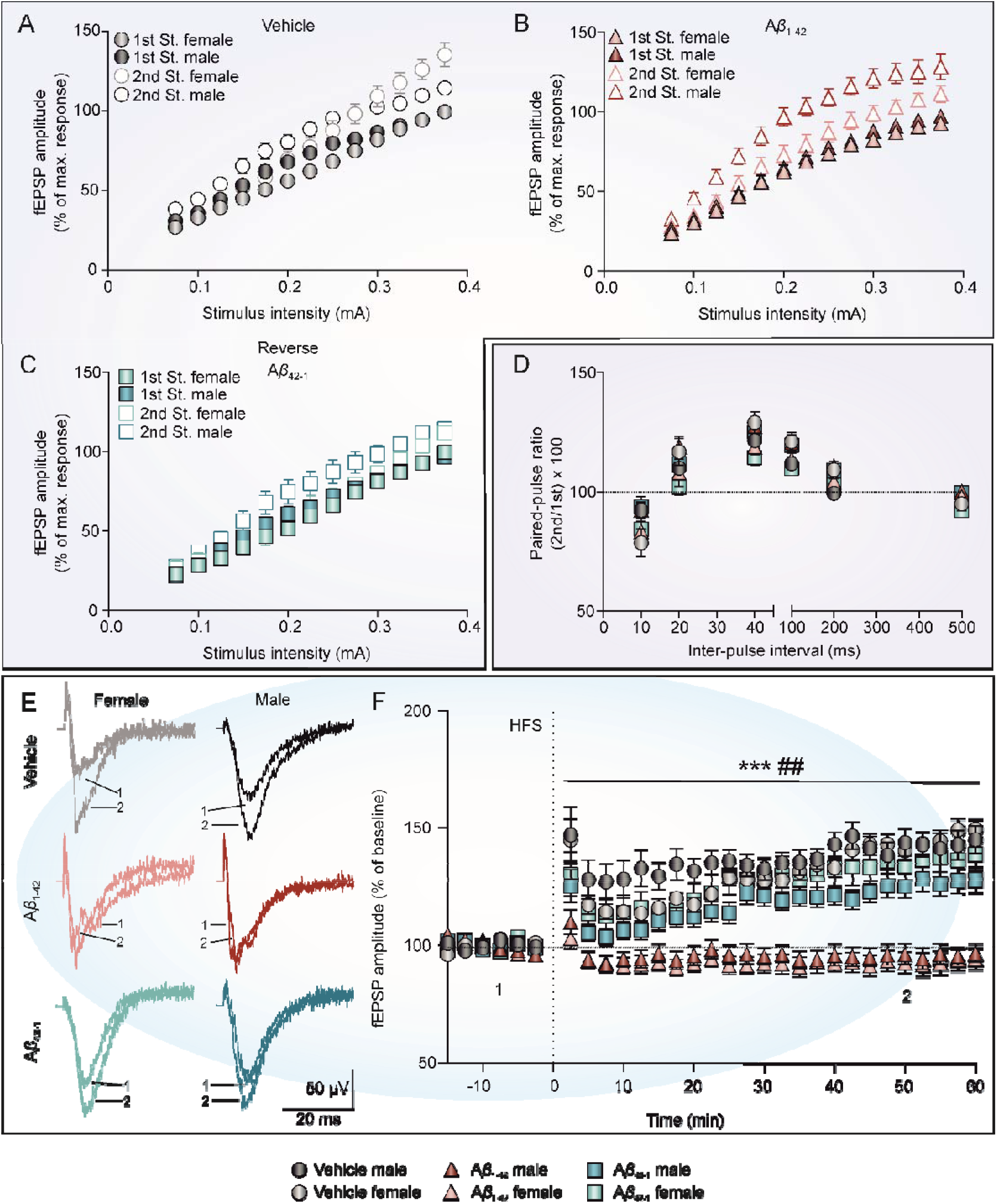
A*β*_1-42_ inhibits ex vivo hippocampal LTP and induces LTD in both female and male mice. **(A-C)** I/O curve with paired fEPSPs collected at increasing stimulus intensities (from 0.075 to 0.4 mA) from control vehicle **(A)**, A*β*_1-42_ **(B)** and A*β*_42-1_ reverse control **(C)** slices, respectively. Data is expressed as a percentage (%) of the maximum amplitude obtained. **(D)** PPF curve with paired fEPSPs collected at interstimulus intervals of 10, 20, 40, 100, 200 and 500 ms. Data is expressed as mean ± SEM amplitude of the second fEPSP as a percentage of the first [(second/first) x 100] for each inter-pulse interval used. **(E)** Representative averaged (n = 5) traces of fEPSPs recorded in the CA1 area, collected during the baseline (1) and ≈50 min post-HFS (2) in hippocampal slices from the different groups. Left traces are from female mice and right traces from male mice. **(F)** Time course of LTP evoked in the CA1 area after HFS in hippocampal slice from the different groups. Recordings were obtained from day 1 to 17 post-*icv.* injection. A*β*, amyloid-*β*; HFS, High frequency stimulation; LTP, long-term potentiation; mA, milliamperes; ms, milliseconds; min, minutes. *** *p* < 0.001 *vs*. vehicle of the corresponding sex; ## *p* < 0.01 *vs*. A*β*_42-1_ of the corresponding sex.

Then, we addressed the short-term plasticity phenomenon, PPF. This protocol is also related to neurotransmitter release and, therefore, allowed us to evaluate the presynaptic functionality after A*β*_1-42_ injection. As shown in Figure 5F, all groups presented an increased response to the second pulse when the intervals were short (20, 40 and 100 ms) since the ratio between the second and first EPSPs were above 100%. Nonetheless, treatment caused no significant differences at any of the selected intervals (vehicle male, n = 6; vehicle female, n = 6; A*β*_1-42_ male, n = 7; A*β*_1-42_ female, n = 5; A*β*_42-1_ male, n = 3; A*β*_42-1_ female, n = 3; Male: F_(2,44)_ = 2.930, *p* = 0.0639; Female: F_(2,39)_ = 1.159, *p* = 0.3243). This data indicated a normal short-term plasticity and presynaptic vesicle release after treatment, suggesting that alterations caused by A*β*_1-42_ injection could be preferentially acting at a postsynaptic level.

Finally, we measured the effect of A*β*_1-42_ injection on long-term synaptic plasticity applying an HFS protocol after a 15-min baseline from day 1 to 17 post- icv. injection (Figure 5G). Data showed a significant treatment effect (Figure 5H; vehicle male, n = 6; vehicle female, n = 9; A*β*_1-42_ male, n = 11; A*β*_1-42_ female, n = 9; A*β*_42-1_ male, n = 10; A*β*_42-1_ female, n = 10; Male: F_(2.78,159.2)_ = 22.87, *p* < 0.001; Female: F_(2,81)_ = 17.67, *p* < 0.001). *Post-hoc* analysis revealed that the differences were specifically between the A*β*_1-42_ and the two control groups, regardless the sex, which showed an inhibited LTP, right after the HFS and during at least 60 min following it. In fact, fEPSPs post-HFS were slightly under the BL for this group, suggesting that a protocol aimed to cause LTP was inducing an LTD instead. As it happened in the memory assays, the control reverse peptide did not affect LTP, implying the specificity of A*β*_1-42_.

Thus, this data indicates that A*β*_1-42_ injection alters long-term synaptic plasticity, mainly at a postsynaptic level, which could be underlying the hippocampal-dependent learning and memory deficits observed in our study in both male and female animals.

## Discussion

The deleterious effect of A*β* - one of the main neuropathological features of AD -, on hippocampal-dependent learning and memory has been widely reported (14). However, the possible differences associated to sex are scarcely explored, despite numerous clinical and epidemiological studies have shown greater prevalence, risk and severity of AD in women (32). Moreover, animal studies, mainly carried out with transgenic mice models, have reported contradictory results (34, 37). Here, we aimed to elucidate whether *icv.* administration of A*β*_1-42_, a local model to study early hippocampal amyloidosis (49, 50), had a differential effect on hippocampal function in female and male mice.

Spatial navigation deficit is an early indicator of AD (51). Furthermore, it has been used to differentiate between various types of dementia, i.e., AD and frontotemporal dementia (52), since AD shows specific deficits in spatial working memory compared to others (53). Here, we used the Barnes maze to assess spatial working, short-term and long-term memory, and the OF habituation test to evaluate a non-associative hippocampal-dependent memory process. Our results showed a similar impairment in both male and female mice in all forms of spatial learning and memory evaluated as well as in the exploratory habituation. It is well stablished that learning and memory deterioration occurs in both, AD patients and animal models (24, 54). Notwithstanding, transgenic models mostly reflect genetic forms of AD and require months to develop the cognitive impairments, whereas AD is mainly a sporadic disorder (55). Moreover, it has been highlighted that AD’s prodromal period lasts from 5 years to even decades (25), with the first cognitive symptom appearing as early as 12 years before dementia onset (56), so it is greatly important to study early stages of the pathology. In agreement with that, A*β* *icv.* administration mimics the sporadic form of AD during early stages. Previous works from our group and others have shown that, even the early amyloidosis caused by a single A*β* injection is sufficient to impair short- and long-term spatial and habituation memory in male rodents (21–23, 29, 30, 53, 57). Moreover, the present study proves that working spatial memory is also altered, as it has been shown in transgenic mice models at later stages of the disease (20, 35). However, this is of special interest in early amyloidosis since working memory deficits can be used to predict which patients with mild cognitive impairment (MCI), a prodromal stage of AD, will eventually develop this type of dementia and the severity of the cognitive decline (58, 59). What is more, the dorsal hippocampus area is implicated in spatial processing, while the ventral hippocampus is related to anxiety-related behaviors (28, 60). Thus, our data show an impairment in spatial learning and memory without affecting anxiety and depression-like behaviors, proving once again that the *icv.* administration of A*β* affect mainly the dorsal hippocampus area (22).

On the other hand, our data did not show any difference between female and male mice neither in the Barnes maze nor in the OF habituation test. Some authors had described similar results, showing no sex-dependent differences in transgenic models (37, 38). Other authors, however, had described the opposite, reporting that females showed more spatial memory deficits than age-matched males also in transgenic mice (33, 35). Nonetheless, those mice displayed prominent amyloid plaques as well as neurofibrillary tangles (34) while in our model of early amyloidosis, A*β* is in its soluble form and not forming senile plaques yet. Thus, it could be hypothesized that in early stages of amyloidosis there is not a sex-dependent alteration, whereas in older animals, with a clearer developed AD pathogenesis, sex becomes an important factor of disease severity. In this line, 3xTg-AD male and female mice, a widely used model of AD, displayed no memory differences at 4 to 10 months of age, but female mice performed worse than males at older ages (20, 61, 62). Interestingly, young females seem to be protected against A*β* toxicity by estrogen, as it promotes a non-amyloidogenic metabolism of APP and has anti-inflammatory properties (32). Later on, when females reach menopause, and therefore estrogen levels decline, this protection is lost, which might underlie the greater susceptibility of women to suffer AD (33). In fact, human studies have reported a negative correlation between estrogen levels and spatial cognition (63). Indeed, estrogen treatment in a menopause-induced 5xFAD mice (another AD mice model) decreases APP and phosphorylated *tau* levels, compared to untreated 5xFAD females (64).

Regarding activity-dependent synaptic plasticity mechanisms underlying learning and memory, our data indicates that a single A*β*_1-42_ injection does not alter short-term plasticity nor presynaptic vesicle release but does affect long-term plasticity at postsynaptic level. Several studies concur with the fact that AD pathology affects mainly at postsynaptic level (65). Early deficits in LTP are associated with the accumulation of A*β* in the hippocampus of transgenic models, and this weakening in synaptic strength is associated with an inability to use cues in a spatial learning task (66). The characteristic hyperexcitability caused by A*β* in the dorsal hippocampus might be increasing the threshold of LTP induction which, following the Bienenstock, Cooper and Munro (BCM) theory, would provoke the induction of LTD since the stimulation fails to properly activate postsynaptic neurons (67). Moreover, A*β* causes an indirect partial unblocking of the synaptic NMDA receptors (68), known to make an essential contribution to spatial working memory processing (69), and activates metabotropic glutamate receptors (70), which result in increased internalization of AMPA receptors, shifting the signaling cascades toward pathways involved in the induction of LTD and synaptic loss (29, 71, 72). This, along with the BCM theory, could partially explain that a protocol aimed to cause LTP induced LTD instead in our female and male A*β*_1-42_ treated animals. In this line, previous work also showed an HFS-induced LTD both *in vivo* (23) and *ex vivo* (22, 53) in male mice after A*β*_1-42_ *icv.* injection. This unbalance in hippocampal synaptic plasticity processes is underlying the hippocampal-dependent learning and memory deficits observed both in male and female animals. Interestingly, LTD, which seems to be increased in AD, is mainly associated with habituation to a novel environment (73), a non-associative type of learning showed to be altered in our mice. This shift in the excitability threshold for LTP/LTD induction caused by A*β* has been shown to be partially governed by G-protein-gated inwardly rectifying potassium (GIRK) channels (22, 43), which main role in the hippocampus is to maintain the inhibitory tone. Indeed, it has been found that their activation with the selective opener ML297 is able to rescue LTP and associated learning and memory processes in this model of early AD amyloidosis (22, 23, 30). Additionally, GIRK2 subunit expression is down-regulated in different models of AD (74, 75) and training in an hippocampal dependent task normalizes its protein level (75). Once again, no differences between male and female mice were found, as expected since there were no sex-dependent changes in memory assessments. Remarkably, our synaptic plasticity results correlate with our behavioral memory experiments, as LTP impairment lasted the same time as memory deficits were observed (i.e., up to 17 days after *icv.* injection) which highlights the importance of this A*β* *icv.* administration model to study early acute-induced AD up to 2 weeks (49, 50). In this line, other authors have shown a progressive degeneration after A*β*_1-42_ intrahippocampal administration that last up to a month (76–78), and *icv*. administration of A*β* mainly diffuses to the dorsal hippocampus (22).

### Perspectives and significance

Despite the growing number of publications studying the sex-related differences underlying the pathogenesis of AD, data regarding females remains inconsistent. In this regard, our results provide systematic evidence of similar hippocampal deficits caused by A*β*_1-42_ regardless the sex, validating this murine model of early amyloidosis. This work opens numerous research perspectives for the future study of AD amyloid-related pathogenesis and treatment in both female and male subjects.

## Conclusions

In summary, our results indicate that a single A*β*_1-42_ *icv*. injection leads to similar and robust habituation and spatial working, short- and long-term memory impairments as well as paired LTP inhibition and LTD facilitation in both sexes. Furthermore, the cognitive and synaptic alterations were long-lasting (observed up to 17 days after treatment), evidencing the convenience of using this *in vivo* mouse model to study early acute stages of amyloidosis regardless the experimental subject’s sex.

## Declarations

### Ethics approval

All experimental procedures were reviewed and approved by the Ethical Committee for Use of Laboratory Animals of the University of Castilla-La Mancha (PR-2022-11-04 and PR-2018-05-11) and conducted according to the European Union guidelines (2010/63/EU) and the Spanish regulations for the use of laboratory animals in chronic experiments (RD 53/2013 on the care of experimental animals: BOE 08/02/2013).

### Consent for publication

Not applicable.

### Availability of data and materials

The datasets used and/or analyzed during the current study are available from the corresponding author on reasonable request.

### Authors’ contributions

LJD and JDNL were responsible for the initial conceptualization; RJH, AC, GIL and DJ performed the experiments; RJH, AC, GIL and SD analyzed the data; RJH and AC were responsible for writing the original draft; RJH, AC, SD, JDNL and LJD did the writing – review and editing. LJD and JDNL were responsible for funding acquisition. All authors read and approved the final manuscript.

## Acknowledgements

Not applicable.

